# Cryo-EM structure of the Smc5/6 holo-complex

**DOI:** 10.1101/2021.11.25.470006

**Authors:** Stephen T. Hallett, Isabella Campbell Harry, Pascale Schellenberger, Lihong Zhou, Nora B. Cronin, Jonathan Baxter, Thomas J. Etheridge, Johanne M. Murray, Antony W. Oliver

## Abstract

The Smc5/6 complex plays an essential role in the resolution of recombination intermediates formed during mitosis or meiosis, or as a result of the cellular response to replication stress. It also functions as a restriction factor preventing viral integration. Here, we report the cryo-EM structure of the six-subunit budding yeast Smc5/6 holo-complex, reconstituted from recombinant proteins expressed in insect cells – providing a full overview of the complex in its apo / non-liganded form, and revealing how the Nse1/3/4 subcomplex binds to the hetero-dimeric SMC protein core. In addition, we demonstrate that a region within the head domain of Smc5, equivalent to the ‘W-loop’ of Smc4 or ‘F-loop’ of Smc1, mediates an essential interaction with Nse1. Taken together, these data confirm a degree of functional equivalence between the structurally unrelated KITE and HAWK accessory subunits associated with SMC complexes.

## INTRODUCTION

The eukaryotic Structural Maintenance of Chromosomes (SMC) family includes the protein complexes cohesin, condensin and Smc5/6. At their respective ‘hearts’ sits an obligate heterodimer of two SMC proteins, either Smc1/Smc3, Smc2/Smc4 or Smc5/Smc6. Globular entities found at both the N- and C-termini of each SMC protein are brought together in space, to form a so-called ‘head’ domain that is capable of binding to, and turning over, ATP. The two halves of the ATPase are connected by a structural excursion known as the ‘arm’; formed from sequential alpha-helical regions that coalesce to form a highly extended anti-parallel coiled-coil. The arm is interrupted at its apex (most distant point from the head) by the ‘hinge’ domain; a region of SMC proteins responsible for hetero-dimerisation with their obligate binding partner. The binding, hydrolysis, and release of ATP by the two head domains (one from each SMC protein) provides a secondary, more transient, and regulated dimerisation interface. The ‘core’ of each SMC complex is then elaborated through binding of additional ‘non-SMC’ protein subunits or ‘elements’, to provide the distinct functionalities required for their respective cellular functions.

For more expansive reviews of the SMC-family, including their respective functions and subunit compositions, please see: (Cutts and Vannini 2020; Datta, Lecomte, and Haering 2020; Hassler, Shaltiel, and Haering 2018; Matityahu and Onn 2021; Palecek 2018; Sole-Soler and Torres-Rosell 2020; Uhlmann 2016; Yatskevich, Rhodes, and Nasmyth 2019).

All three complexes are required for the organisation and management of chromosome architecture and structure throughout the cell cycle. Cohesin has well described roles in sister chromatid cohesion and the organisation of the interphase chromosomes into topologically associated domains or TADs, whereas condensin is required to compact chromosomes at mitosis. The Smc5/6 complex has roles in the processes of DNA replication and DNA damage repair; the complex acting to suppress / prevent formation of inappropriate structures that can form during homologous recombination-mediated rescue of replication forks that have stalled (or collapsed) on encountering replication ‘road-blocks’ or obstacles.

Alterations to the coding sequence of human Nse2 (Non-SMC-element; generally written as either Nse or NSMCE) have been linked to primordial dwarfism and insulin resistance (Payne et al. 2014), with changes in Nse3 linked to severe lung disease immunodeficiency and chromosome breakage syndrome (LICS, van der Crabben et al. 2016). Interestingly, Smc5/6 is also specifically targeted for ubiquitylation and degradation by the regulatory protein X of hepatitis B virus (HbX) to alleviate restriction of viral replication by the complex (Murphy et al. 2016). The complex has also been shown to act as a restriction factor working against other viruses (Bentley et al. 2018; Gibson and Androphy 2020; Xu et al. 2018).

As well as containing subunits that provide both ubiquitin and SUMO E3-ligase activity (through Nse1 and Nse2 respectively), Smc5/6 is further differentiated from the cohesin and condensin complexes by the fact that the two non-SMC proteins (Nse1 and Nse3) that bind to its kleisin subunit (Nse4) belong to the ‘KITE’ family kleisin-interacting tandem winged-helix element; Palecek and Gruber 2015) rather than the distinct and structurally unrelated ‘HAWK’ family (HEAT proteins associated with kleisins; Wells et al. 2017). As proteins of the KITE-family are also found in prokaryotic SMC complexes, it has led to the hypothesis that Smc5/6 best represents the eukaryotic ‘cousin’ of *Bacillus subtilis* Smc/ScpAB and *Escherichia coli* MukBEF (Palecek and Gruber 2015) — however, it is still unclear as to when in the evolutionary timescale KITEs were replaced by HAWK subunits, to create cohesin and condensin complexes.

Here, we present the cryo-EM structure of the budding yeast Smc5/6 complex, in its apo or non-liganded form. Our study serves to confirm the overall architecture of the complex, plus provide details of how and where the Nse1/Nse3/Nse4 KITE-kleisin subcomplex interacts with the Smc5/Smc6/Nse2 core. Additional experiments reveal the presence of a crucial interface formed between the head domain of Smc5 and Nse1, which utilises the equivalent of the ‘W-loop’ or ‘F-loop’ found in Smc4 and Smc1 respectively (Hassler et al. 2019). Taken together, our data uncover an unanticipated degree of functional equivalence between KITE and HAWK accessory subunits.

## RESULTS

In this manuscript, for simplicity and brevity, we refer to the protein components of the *Saccharomyces cerevisiae* SMC complexes, unless otherwise indicated.

We have previously described reconstitution and characterisation of the *S. cerevisiae* Smc5/6 complex, using recombinant proteins expressed in insect cells (Hallett et al. 2021). Here, cryo-EM data were collected for the six component ‘holo-complex’, where Smc5 and Smc6 contain Walker B mutations E1015Q and E1048Q, respectively (**Fig. 1a, Materials and Methods**).

**Fig. 1.**
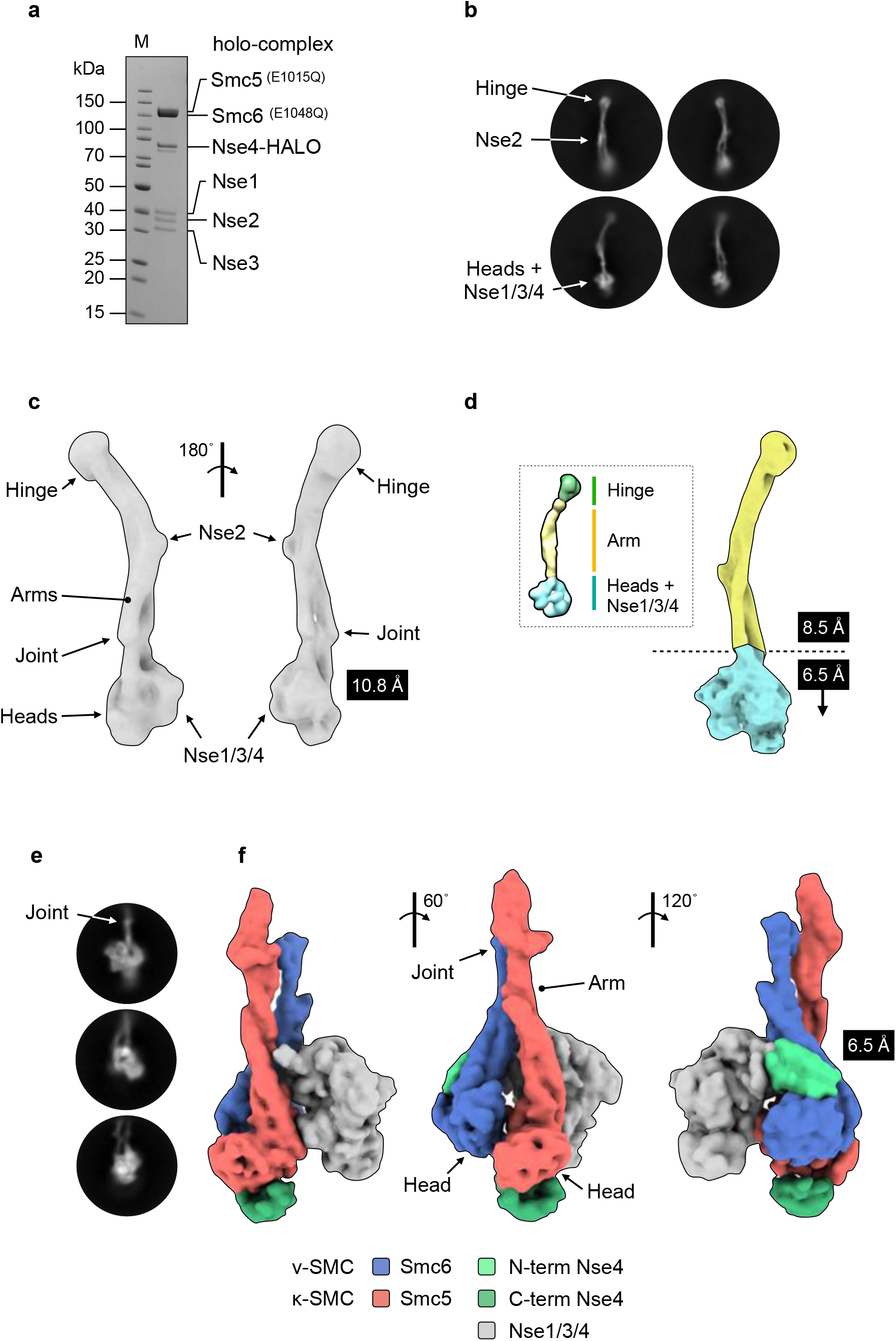
Cryo-EM of the budding yeast Smc5/6 complex. (a) Representative SDS-PAGE gel for the purified Smc5/6 holo-complex. (b) Representative 2D class averages (side views). Conformational flexibility leads to blurring of density at either the head (top) or hinge-end (bottom) of the complex. (c) Initial 3D map from cryo-EM at a resolution of 10.8 Å. (d) Maps from focussed refinement at 8.5Å and 6.5 Å for the indicated segments of the Smc5/6 holo-complex (d, inset) model obtained by uranyl acetate negative stain electron microscopy for comparison (Hallett et al. 2021). (e) Representative 2D class averages (side views) and (f) resultant 3D map at 6.5 Å for the ‘head’-end of the complex. The cryo-EM map has been segmented and coloured with respect to its assigned component (see associated key for additional detail).

A neural network was trained with a set of manually picked particles (Bepler et al. 2019). The resultant ‘picking model’ was then used to identify a total of 380,714 particles (see **Sup. Fig. 1** for data-processing summary). Two sets of two-dimensional (2D) class averages emerged from processed data, with either the ‘hinge’ (**Fig. 1b, top**) or the ‘head-end’ of the complex more clearly in focus (**Fig. 1b, bottom**); the blurred density at either end consistent with a degree of conformational flexibility within the coiled coil ‘arms’ (Hallett et al. 2021). From this, a sub-set of 17,162 particles were used to generate a medium-resolution 3-dimensional (3D) map that covered the entire length of the holo-complex, reconstructed at a resolution of 10.8 Å as judged by the 0.143 Fourier shell correlation (FCS) criterion (Rosenthal and Henderson 2003) (**Fig. 1c**). Focussed refinement allowed a slightly higher resolution map to be calculated for the upper (106,660 particles, 8.5 Å) segment of the complex (**Fig. 1d**). Pleasingly, resultant maps were consistent with the envelope we previously obtained by uranyl acetate negative stain transmission electron microscopy (Hallett et al. 2021), **Fig. 1d, inset**).

In parallel, we trained a second neural network with manually picked particles that encompassed just the head-end of the complex (824,644 particles). The resultant 2D class averages and 3D map from processed data (84,180 particles, 6.5 Å) are shown in **Fig. 1e** and **1f** respectively. Segmentation analysis (Pettersen et al. 2021; Pintilie and Chiu 2012; Pintilie et al. 2010) allowed portions of the map to be readily assigned to the head domains, as well as the Nse1/Nse3/Nse4 (Nse1/3/4) subcomplex (coloured red, blue and grey respectively in **Fig. 1f**). Additional segments of density associated with either the ‘head’ or ‘arm’ (dark or light green, **Fig. 1f**) allowed the identity of each head-domain to be determined, due the expected binding positions of the N- and C-terminal domains of the kleisin Nse4 (please see additional text below).

### A model for the Smc5/6 holo-complex

A sharpened composite map, generated from the highest resolution maps for each segment of the holo-complex, was of sufficient quality to allow positioning of the coiled-coil regions for both Smc5 and Smc6, as well as placement of the structure of *S. cerevisiae* Nse2 (Mms21) bound to the arm of Smc5 (PDB: 3HTX; Duan et al. 2009). This, combined with homology models for the ‘hinge’, ‘heads’ and Nse1/3/4, allowed an initial pseudo-atomic model to be constructed; this model was subsequently updated using AlphaFold predictions made available through the EMBL-EBI repository (https://alphafold.ebi.ac.uk; Jumper et al. 2021) (see **Materials and Methods**).

Working from the ‘top’ of the complex downwards, we see that the hinge domain is tilted with respect to an axis defined by the coiled-coil arms; consistent with our previous analysis of this region in the fission yeast complex and with data published for condensin (Alt et al. 2017; Soh et al. 2015). Below this, there is an apparent discontinuity / break in the helices of the arms, located between the hinge and Nse2 subunit — a feature likely to represent the ‘elbow’ found in the other complexes of the SMC-family (Burmann et al. 2019; Kong et al. 2020; Lee et al. 2020) — notably, at this point the two arms of Smc5/6 also cross over each other (**Fig. 2a**, inset *ii*).

**Fig. 2.**
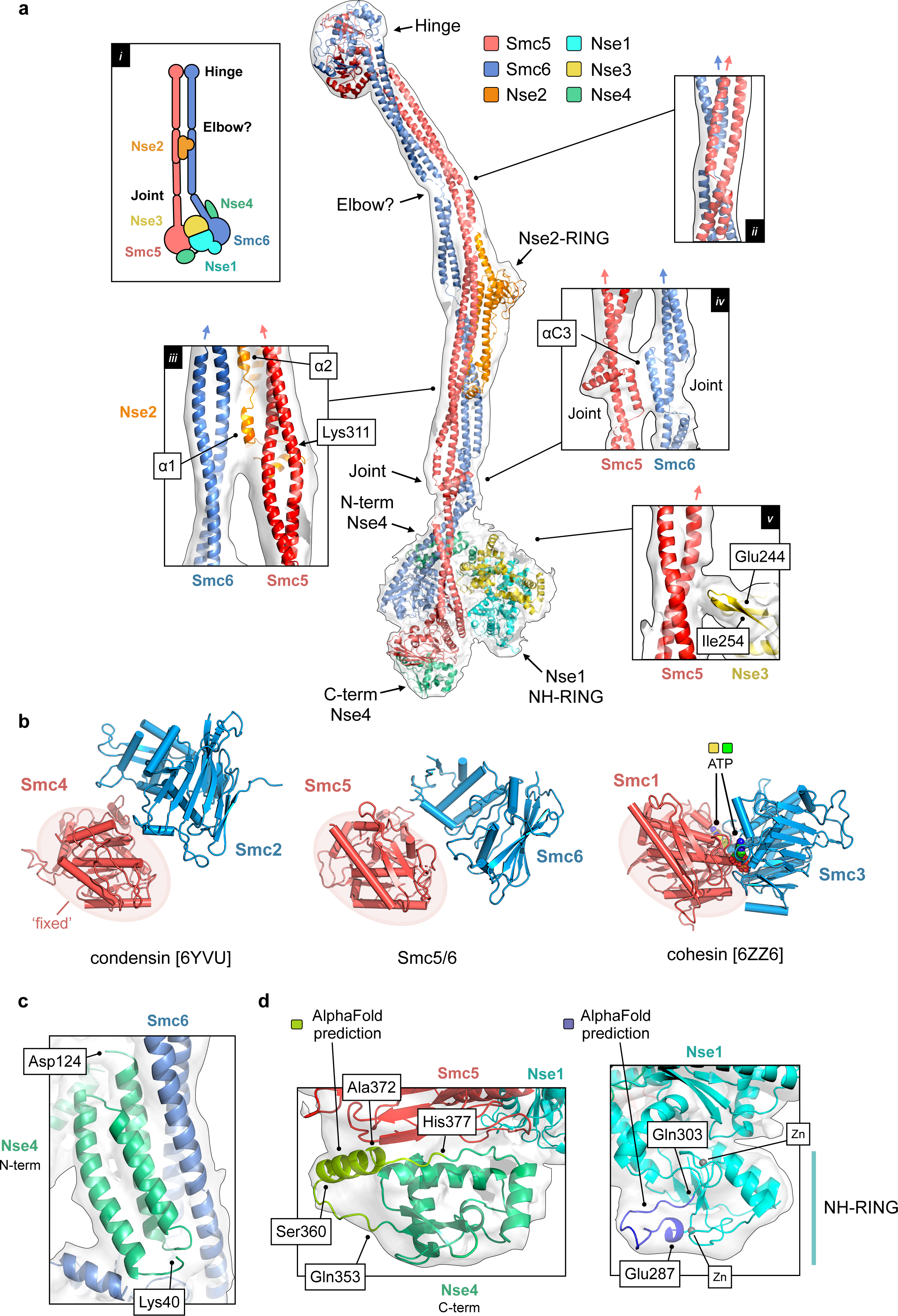
A pseudo-atomic model for the Smc5/6 holo-complex. (a) Overview of the pseudo-atomic model. (a, inset *i*) Schematic showing the overall architecture of the Smc5/6 complex and selected molecular features. (a, inset *ii*) Expanded view of the ‘Elbow’, highlighting the crossover of the coiled-coil ‘arms’ of Smc5 and Smc6 at this point. (a, inset *iii*) The first alpha-helix of Nse2 (α1) is situated between the two arms of the complex. The position of Lys311, a known site of auto-SUMOylation is also indicated (a, inset *iv*) Expanded view of the interface between the ‘Joint’ features of Smc5 and Smc5, involving the two αC3 helices. (a, inset *v*) A short beta-hairpin (amino acids Glu244-Ile254) protruding from Nse3 is in close proximity to the arm of Smc5. For each inset, the directionality of the ascending helix (head to hinge) is indicated by a blue or red arrow, for Smc5 and Smc6 respectively. (b) Comparison of the relative head domain positions in the cryo-EM structures of budding yeast condensin (PDB: 6YVU), Smc5/6 (this manuscript) and cohesin (PDB: 6ZZ6); in each, using the head of the κ-SMC as a fixed reference point. (c) Expanded view for the N-terminal helical domain of Nse4 (aa Lys40-Asp124) bound to the ‘arm’ of Smc6. (d, left) AlphaFold predicts the presence of an additional helical element (aa Ser360-Ala372) in the C-terminal domain of Nse4. (d, right). AlphaFold predicts a budding yeast-specific loop insertion in the NH-RING of Nse1 (aa Glu287-Gln303). Where shown, sections of density from the composite cryo-EM map are represented by a semi-transparent molecular surface, shaded in grey. Please also see associated key for additional detail.

In our previous lower-resolution study, we observed that the arms of Smc5/6 remain in close proximity (‘arms-together’, ‘rod-like’, ‘I’-conformation) for the majority of their length (Hallett et al. 2021), in agreement with parallel studies published by other laboratories (Gutierrez-Escribano et al. 2020; Taschner et al. 2021; Yu et al. 2021). With the additional resolution now afforded by cryo-EM, we see that the arms separate slightly from each other after the junction with Nse2 (**Fig. 2a**, inset *iii*) but then remain approximately parallel until the head-end of the complex. Our model positions the first alpha-helix (α1) of Nse2 such that it can ‘talk’ across to the descending helix (hinge to head) of Smc6, as well as make its previously documented set of interactions with Smc5 (Duan et al. 2009); the helix appearing to ‘glue’ the two arms together, in agreement with crosslinking data for the holo-complex recently published by Taschner et al. (2021) (**Sup. Fig. 2a**). The two arms then briefly re-contact each other, through the two inward facing helices (αC3) of the ‘joint’ (**Fig. 2a**, inset *iv*) — a molecular feature common to SMC proteins (Diebold-Durand et al. 2017) formed from three helix-loop repeats that encircle the more continuous ascending alpha-helix, thus generating both an interruption and point of flexure within the coiled-coiled structure of the arm; again supported by crosslinking data (**Sup. Fig. 2b**; Taschner et al. 2021).

In our model, the head domains of Smc5 and Smc6 are not in direct contact. Comparison of head domain positions indicates that our structure most resembles that of the apo / ATP-free / ‘non-engaged’ / juxtaposed / or ‘J’-state of budding yeast condensin deposited under PDB accession code 6YVU (Lee et al. 2020) rather than the ATP-bound / ‘engaged’ / ‘E’-state seen for budding yeast cohesin when in complex with Scc2 and DNA (PDB: 6ZZ6; Collier et al. 2020) (**Fig. 2b**). Sections of the map corresponding to the RecA lobe of both the Smc5 and Smc6 head domain become less apparent at higher contour levels, suggesting a degree of conformational flexibility in this part of the complex, as supported by estimates of local resolution (**Sup. Fig. 2c**).

### Revealing the binding location of the Nse1/3/4 sub-complex

The Nse1/3/4 subcomplex is located to one side of the central axis defined by the coiled-coil arms of Smc5 and Smc6. The winged-helix 2 (WH/2) domain of Nse1 and the head domain of Smc5 are in direct contact, with a short loop protruding from the equivalent winged-helix domain of Nse3 (WH/B) positioned to interact with the arm of Smc5 (amino acids 244-254, **Fig. 2a** inset *v*). Notably, neither Nse1 nor Nse3 directly interact with Smc6.

The recent publication of an X-ray crystal structure for *Xenopus laevis* Nse1/3/4, has provided molecular details for how the kleisin subunit interacts with the Nse1/Nse3 KITE heterodimer (kleisin-interacting tandem winged-helix element). The central section of Nse4 follows a path through the centre of both KITE proteins, interacting with the linker regions that serve to connect their component winged-helix domains together (Jo et al. 2021). Pleasingly, our cryo-EM data allows integration of this kleisin path with the set of interactions made by the globular domains found at each of its termini: the helical N-terminal domain binding to the ‘neck’ of Smc6 (**Fig. 1f and 2c**) and the C-terminal domain to the ‘cap’ of the Smc5 head domain (**Fig. 1f and 2d**, **left**) thus confirming at the structural level the set of kleisin-facilitated interactions conserved across the SMC-family of complexes, including prokaryotic ScpAB and MukBEF as well as eukaryotic condensin and cohesin (Burmann et al. 2013; Diebold-Durand et al. 2017; Fennell-Fezzie et al. 2005; Gligoris et al. 2014; Haering et al. 2004; Kamada et al. 2017; Woo et al. 2009; Zawadzka et al. 2018). Saying this, the resolution of our composite map is not sufficient to allow an unambiguous tracing of the parts of Nse4 that serve to connect the N-terminal domain to the central section (amino acids 125-183) or the central section to the C-terminal domain (aa 46-283), indicating either intrinsic disorder or a high degree of conformational flexibility; an observation consistent with the kleisin moieties in cryo-EM structures of other eukaryotic SMC complexes (Collier et al. 2020; Higashi et al. 2020; Lee et al. 2020; Shi et al. 2020).

### Additional structural features are predicted by AlphaFold

AlphaFold predicts (albeit with a lower level of confidence) the presence of a hereto unknown alpha-helical element in the C-terminal domain of Nse4 (amino acids Ser360 through to Ala372). Nicely, this provides a facile explanation for a region of additional density visible in our composite map, not accounted for by the initial homology model (**Fig. 2d left**). Similarly, AlphaFold predicts the presence of a budding yeast-specific extended loop within the NH-RING domain of Nse1 (amino acids Glu287 to Gln303), which again provides a better overall fit to the experimental map (**Fig. 2d right, Supp. Fig 2d**).

### A loop in Smc5 has functional equivalence to the W-loop of Smc4 and F-loop of Smc1

The location of Nse1/3/4 binding is particularly striking, as it echoes that of the structurally unrelated HAWK accessory proteins (HEAT proteins associated with kleisins) found in condensin and cohesin complexes; in each case, the head domain of the κ-SMC (Smc5, Smc4, Smc1) serving to provide a major interaction surface (**Fig. 3a**).

**Fig. 3.**
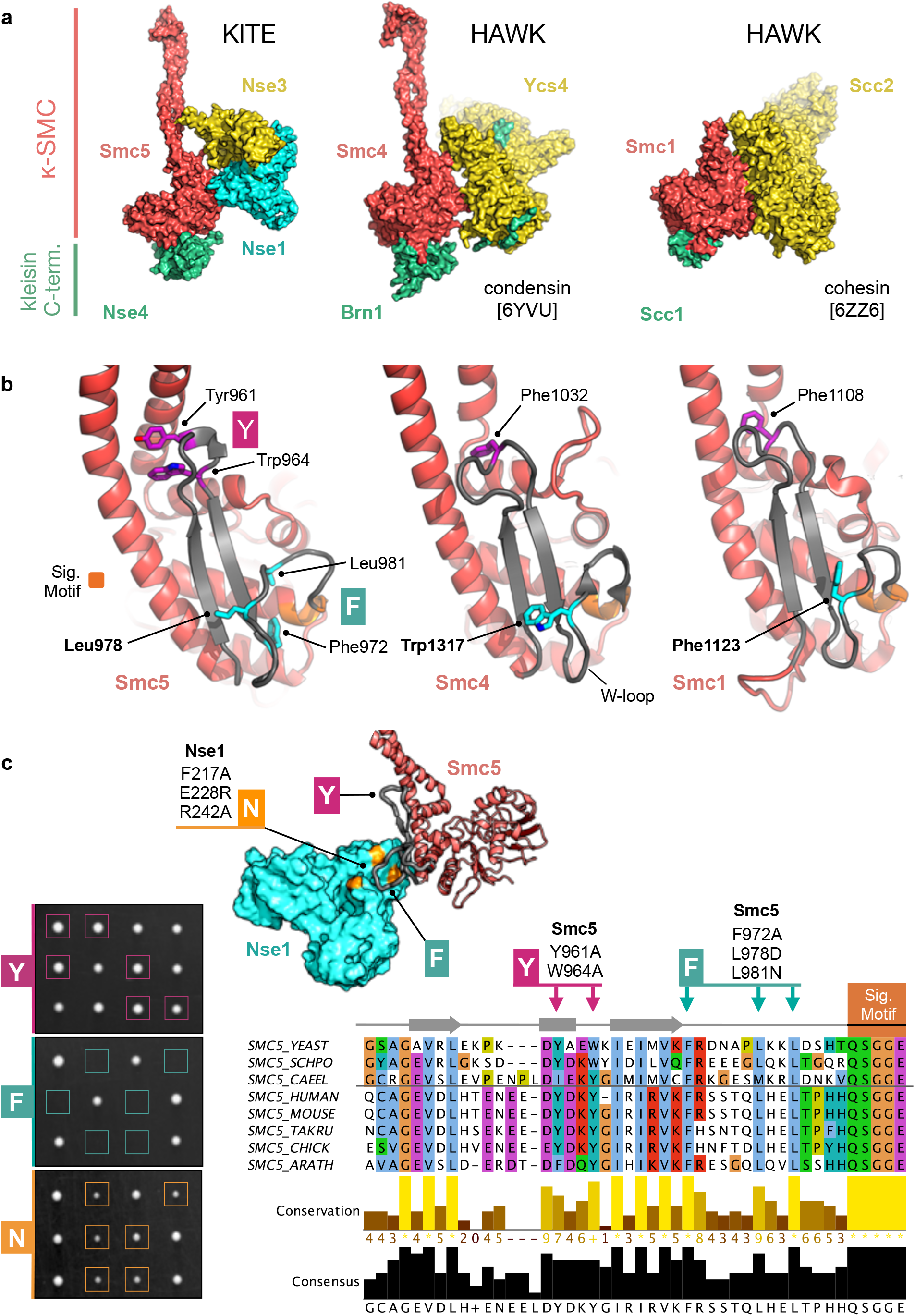
KITES and HAWKS share a common interaction interface that involves the κ-SMC ‘W-loop’. (a) Side-by-side visualisation of the κ-SMC head domain from Smc5/6 (left), condensin (middle) and cohesin (right) in complex with their respective kleisin C-terminal domain; Nse4, Brn1 and Scc1. In each case, the interacting partner, whether KITE or HAWK, makes a similar set of interactions with the head domain of the κ-SMC. (b) Expanded view, showing secondary structure molecular cartoons for each κ-SMC head domain, highlighting the position of conserved amino acids within the ‘W-loop’ or equivalent (stick representation, carbon atoms coloured cyan) plus aromatic residues within the preceding loop (stick representation, carbon atoms coloured magenta). The ABC-signature motif is additionally highlighted in orange. (c, left) Tetrad dissections. Spores derived from diploid *S. cerevisiae* strains carrying both wild-type allele and indicated mutant allele plus associated NAT-selectable marker (natMX6). Genotypes were confirmed by replica plating of spores on selective media (not shown). (c, right) Multiple sequence alignment, across selected species, showing conservation and consensus of amino acids within the W-loop and preceding region of Smc5 (produced using Jalview 2 with Clustal X colour scheme; (Waterhouse et al. 2009)). (c, top left) Relative positions of the Y and F mutation sets, with respect to the head domain of Smc5 and Nse1 subunit (c, inset) Molecular secondary structure cartoon / surface representation, showing the relative locations of the introduced mutation sets within the Nse1/Smc5 interface. Sets of compound mutations introduced into budding yeast: Smc5-Y = Y961A, W964A; Smc5-F = F972A, L978D, L981N, Nse1-N = F217A, E228R, R242A. Please also see associated key for additional detail.

Closer inspection reveals that a region of Smc5 (amino acids Gly947—Gly978), which ‘talks’ across to the Nse1 subunit, is structurally equivalent to the ‘W-loop’ of the Smc4 head domain (Hassler et al. 2019) (**Fig. 3b, 3c inset**). Notably, defined mutations (S1316D or W1317A) introduced into the W-loop render budding yeast cells non-viable. Furthermore, in the context of a fully recombinant ‘head complex’, mutation of the equivalent tryptophan residue in *Chaetomium thermophilum (Ct*) Smc4 resulted in a dramatic reduction of its ability to turnover ATP (Hassler et al. 2019). In Smc1, the equivalent region has been given the alternative title of ‘F-loop’, due to the presence of a conserved phenylalanine residue (F1123 in budding yeast Smc1 (Hassler et al. 2019; Petela et al. 2021; Shi et al. 2020).

In Smc5, Leu978 appears to be the structurally equivalent ‘F’ or ‘W-loop’ residue (**Fig. 3b**) — a relationship supported by a strong preference for a leucine in this position, as revealed by a cross-species multiple amino acid sequence alignment (**Fig. 3c**). However, two additional amino acids within the same loop, Phe972 and Leu981, are also strictly conserved (label F in **Fig. 3b** and **c**). A second region of conservation was also evident in the preceding loop (labelled Y in **Fig. 3b and c**). Here, structural comparison suggested that one or more aromatic residues might potentially act to anchor or connect the helical lobe of the head domain back to its coiled-coil arm. Interestingly, in *E. coli* MukB, this region is highly elaborated to form the so-called ‘larynx’, residues of which contribute to its DNA-binding interface (Bürmann et al, 2021; https://doi.org/10.1101/2021.06.29.450292).

When working with homology models and structural data of limited resolution, there is a degree of uncertainty as to the precise amino acids involved in a molecular interface. With this in mind, we introduced sets of mutations designed to either disrupt localised structure or alter key features within a loop (**Materials and Methods**). Yeast harbouring the (Y) Y961A, W964A double mutation were viable and surprisingly displayed no sensitivity to a range of genotoxic agents. In sharp contrast, introduction of the (F) F972A, L978D, L981N triple mutant rendered budding yeast cell inviable, as determined by dissecting sporulated diploids (**Fig. 3c, Sup. Fig. 3a**); confirming the importance of this loop to cellular function. We also introduced a set of complementary mutations into Nse1, designed to perturb the interaction with the Smc5 loop (N: F217A, E228R, R242A). Here, yeast were viable, but had a slow growth phenotype and mild sensitivity to treatment with MMS or camptothecin (**Fig. 3c and Sup. Figs. 3c and 3d**).

## DISCUSSION

Using recombinant proteins expressed in insect cells we have reconstituted, then visualised by cryo-EM, the six-subunit budding yeast Smc5/6 ‘holo-complex’ — in its apo or unliganded form — to provide a structural overview of the complex’s architecture and reveal the position of the bound Nse1/3/4 sub-complex. Whilst structurally unrelated to the HAWK-family of proteins (HEAT proteins associated with kleisins; Wells et al. 2017) the set and type of interactions made by the KITE (kleisin-interacting tandem winged-helix element; Palecek and Gruber 2015) hetero-dimer of Nse1/3 and the kleisin Nse4 indicate several unifying features that span the SMC-family of complexes:

### i) KITE and HAWK subunits can bind to double-stranded DNA

**Smc5/6:** The ability of the Smc5/6 complex to bind dsDNA is provided by the Nse1/3 heterodimer (see also text below, Zabrady et al. 2016). Furthermore, reconstituted complexes lacking both Nse1/3 and the kleisin Nse4 do not bind dsDNA (Hallett et al. 2021). In condensin and cohesin, a similar dsDNA-binding activity is provided by the HAWK subunits, with the notable exception of Pds5 (Li et al. 2018). **Condensin:** Both the Brn1-Ycg1-Ycs4 hetero-trimer (Piazza et al. 2014) as well as Ycg1 in complex with the C-terminal portion of Brn1 (Kschonsak et al. 2017) can interact with nucleic acid. Whilst DNA-binding by Ycs4 has not been formally demonstrated, an examination of its structure and surface charge suggests the presence of a compatible region located between HEAT repeats 7 and 8 (Lee et al. 2020); a view supported by experiments using *Ct*Ycs4 and *Ct*Ycg1 which, as isolated proteins, have affinity for dsDNA (Piazza et al. 2014). **Cohesin:** Scc3, when bound to Scc1, is capable of binding to dsDNA (Li et al. 2018). DNA-binding has been reported for Mis4, the *S. pombe* equivalent of Scc2, both in the context of its hetero-dimeric complex with Ssl3 (equivalent to Scc4) or as an isolated protein (Murayama and Uhlmann 2014; Kurokawa and Murayama 2020). Recent data indicates a similar pattern of behaviour for both the Scc2/Scc4 hetero-dimer and for Scc2 alone [Collier *et al*., 2020; https://doi.org/10.1101/2020.06.03.132233].

### ii) Addition of dsDNA stimulates ATPase activity

**Smc5/6**: Addition of dsDNA to Smc5/6 strongly stimulates its ATPase activity (Fousteri and Lehmann 2000) with enhancement being strictly dependent on the presence of the Nse1/3/4 sub-complex (Hallett et al. 2021). **Condensin:** Similarly, the Brn1-Ycg1-Ycs4 or ‘non-SMC’ complex is essential for dsDNA-dependent simulation of ATPase activity (Piazza et al. 2014). **Cohesin:** Here, the Scc2 subunit is required, as determined by experiments that added the protein to pre-assembled complexes containing either the core hetero-trimer (Smc1/Smc3/Scc1) or hetero-tetramer (Smc1/Smc3/Scc1/Scc3) (Petela et al. 2018). A similar requirement has been demonstrated for the fission yeast cohesin complex (Murayama and Uhlmann 2014). In each case, the presence of the kleisin subunit is required to connect the dsDNA binding activity of the KITE/HAWK protein to the core SMC complex.

### iii) dsDNA-binding is intrinsically linked to chromatin-binding, loading or retention

**Smc5/6:** Hypomorphic (non-lethal) mutations that reduce the ability of the Smc5/6 complex to bind to dsDNA, result in a loss of chromatin association, as shown by chromatin immuno-precipitation (ChIP) and live-cell single-molecule tracking experiments (Etheridge et al. 2021; Zabrady et al. 2016). **Condensin:** Mutations that reduce the ability of Brn1 to interact with dsDNA, as part of its composite interaction surface with Ycg1, reduce the amount of condensin associated with chromatin at defined loci (Kschonsak et al. 2017). **Cohesin:** Mutations that affect the ability of Scc2 or Mis4 to bind DNA result in reduced chromatin binding/loading, at several known cohesin-enriched loci (Collier et al. 2020; Kurokawa and Murayama 2020).

### iv) mutations within the W-loop (or equivalent) of the κ-SMC perturb function

The ‘W-loop’ is a conserved structural feature, present in each of the κ-SMC subunits (Smc1, Smc4, Smc5) albeit with different patterns of amino acid conservation. Mutation of highly conserved amino acids within the loop are incompatible with cell viability of yeast when introduced into Smc4 (Hassler et al. 2019) or Smc5 (this manuscript).

By taking each of the above points in hand, we can now add the Nse1/3 hetero-dimer into the statement made by Petela et al. (2018) “A role in contacting DNA might therefore be a feature conserved between Scc2 and Ycg1” and also amend the statement made by Hassler et al. (2019) that ‘…interaction of a HEAT-repeat subunit with the κ-SMC_hd_ (head domain) is a central feature of all condensin and cohesin complexes’ — to also include Smc5/6 — to state that ‘interaction of a kleisin-associated subunit, whether HAWK or KITE, with the κ-SMC head domain is a central feature of the hetero-dimeric SMC complexes, condensin, cohesin and Smc5/6.’

### A speculative model for DNA-binding and engagement

Zabrady et al. (2016) previously reported a docking pose for binding of a double-stranded DNA duplex to a positively charged cleft formed between human Nse1 and Nse3 (NSMCE1/NSMCE3) supported by a range of biochemical and biophysical data. A more recent study using *Xenopus laevis* Nse1/3/4 has reported a similar mode of interaction (Jo et al. 2021). By simple superposition, one end of the docked DNA duplex would physically clash with the arm of Smc5 (**Fig. 4a, left**). However, the DNA duplex could be accommodated by a relatively small movement of the Nse1/3/4 subcomplex away from the Smc5 head domain (or vice versa). Saying this, the resulting pose would still be at odds with the expected dsDNA binding location, as illustrated here by comparison to the DNA-bound *S. cerevisiae* cohesin complex (**Fig. 4a, right**. SMC1-SMC3-SCC1-SCC2 complex; PDB: 6ZZ6), but also observed for other proteins of the SMC-family, including the more distantly related Rad50 (Kashammer et al. 2019).

**Fig. 4.**
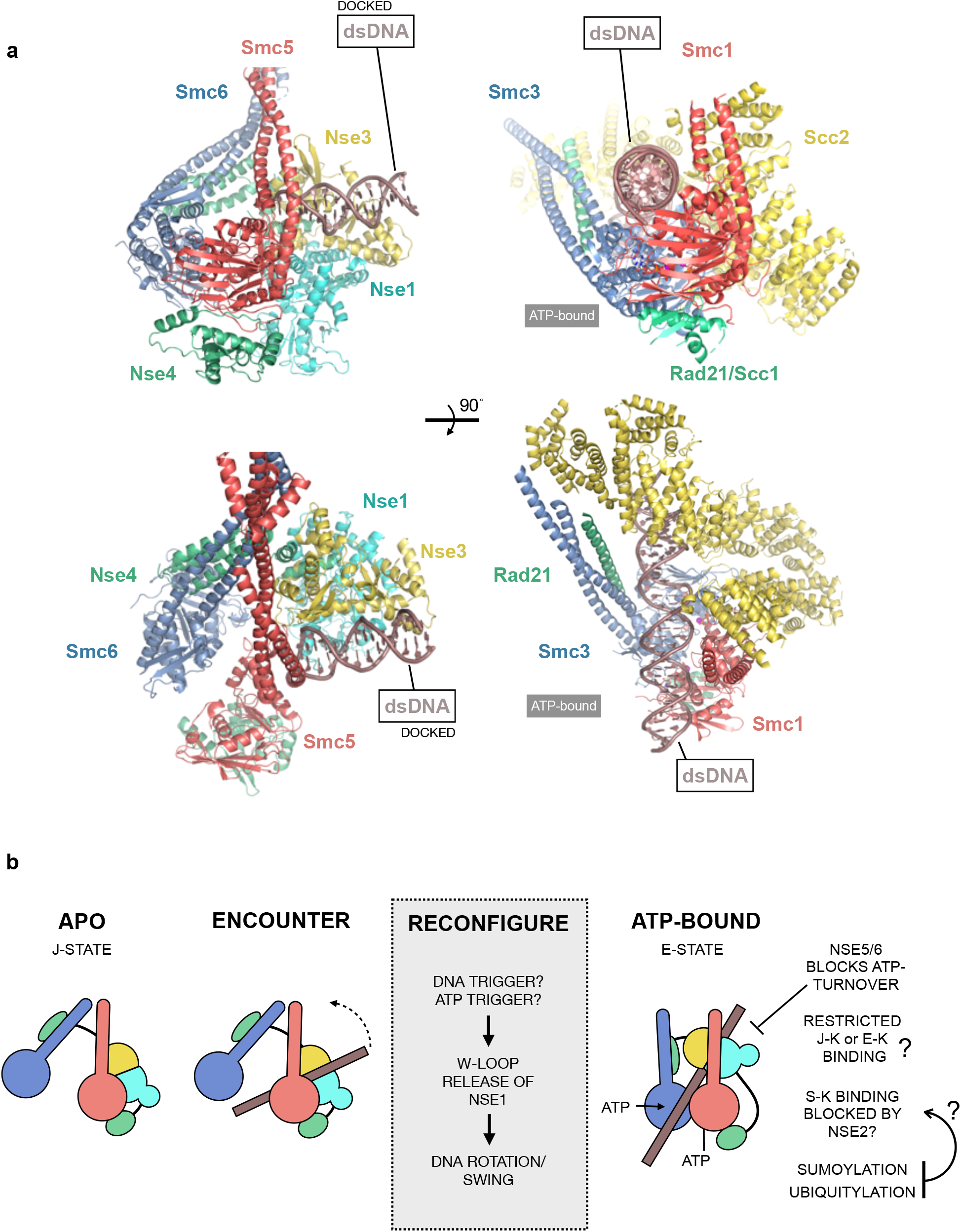
A speculative model for dsDNA binding by the Smc5/6 complex. (a) Secondary structure molecular cartoons comparing a docked pose for dsDNA-binding to the Smc5/6 complex with cohesin in complex with ATP and dsDNA (PDB: 6ZZ6). The trajectory of bound DNA was obtained by superposition of the docked pose reported in (Zabrady et al. 2016) for the human NSE1/3 hetero-dimer. (b) Model for conformational changes upon DNA binding. We propose that transition from the apo, non-engaged state (J-state) to ATP-bound, engaged (E-state) includes an intermediary ‘encounter’ complex — where dsDNA is first bound to a positively charged surface/groove (created by the interface between Nse1 and Nse3) — before a secondary reconfiguration step, triggered by either DNA or ATP-binding, acts to release the Smc5 ‘W-loop’ from its interaction with Nse1. This allows stimulation of ATP turnover by the head domains and repositions the dsDNA such that it is bound with the expected configuration (i.e., as in cohesin). It is not clear how, or indeed if, ubiquitylation, SUMOylation or other post-translational modification affects either conformation or ATPase activity. It is also not known if the presence of Nse2, acts to block binding or transition of bound dsDNA into the S-K ring (SMC-kleisin) compartment, or if it serves to restrict it to the J-K / E-K (juxtaposed-heads/kleisin or engaged head-kleisin) spaces. Binding of the Nse5/6 heterodimer blocks the ability of Smc5/6 to turn over ATP (Hallett et al. 2021; Taschner et al. 2021), but it is not known what effect this has on the overall conformation at the head-end of the complex.

In their 2019 paper, Hassler et al. state that ‘It is likely that tilting and repositioning of the W-loop residues dissociates Ycs4, which binds this part of Smc4_hd_ (head-domain) and sterically blocks access of Smc2_hd_’. The follow-on study builds on this initial hypothesis to uncover a ‘flip-flop’ mechanism, which involves a physical switch of HEAT subunit from Ycs4 (binding to the head domain of Smc4) in the apo/unliganded state, to Ycs1 (binding to the head domain of Smc2) in the engaged/ATP-bound state (Lee et al. 2020). It is therefore tempting to speculate that a similar mechanism may well be at play in Smc5/6, as in order for the head-domain of Smc5 to become engaged with that of Smc6, a conformational change is dictated — potentially linked to breaking of the Smc5 ‘W-loop’ / Nse1 interface, to release and allow repositioning of the KITE sub-complex, and thus facilitate engagement of a bound dsDNA substrate with the top section of each head domain (see speculative model presented in **Fig. 4b**). Here, however, a ‘flip-flop’ mechanism might invoke the Nse3 subunit and a potential direct interaction with the head domain of Smc6. This will, however, require experimental validation, especially given that in PDB entry 6ZZ6 (cohesin) the F-loop of Smc1 remains engaged with the Scc2 subunit, indicating that the precise molecular choreography underpinning the initial ‘engagement’ complex is still unclear, and may well be (subtly?) different for each complex of the SMC-family in accordance with their respective cellular functions.

### Does Nse2 prevent or control bending at the elbow?

We and others have observed some degree of flexure at the ‘elbow’ of Smc5/6 (Gutierrez-Escribano et al. 2020; Hallett et al. 2021; Serrano et al. 2020; Taschner et al. 2021; Yu et al. 2021) but without the acute bend observed in condensin, cohesin and MukBEF (Anderson et al. 2002; Burmann et al. 2019; Collier et al. 2020; Higashi et al. 2020; Kong et al. 2020; Lee et al. 2020; Shi et al. 2020). However, the set of interactions made by Nse2 with Smc5, and now Smc6, may provide insight, as these are likely to prevent separation and rotation of the arms, and therefore the concerted set of motions thought to produce the bend (Burmann et al., 2019).

Interestingly, a major hotspot for auto-SUMOylation has been mapped to a region within the coiled-coil arm of Smc5, sitting between amino acids Lys310 and Lys327. Whilst the precise site(s) of modification appears to be redundant, Lys311 has been identified with high confidence as a major site of auto-SUMOylation (Zapatka et al. 2019). Saying this, it is not known what effect SUMOylation has on the overall structure / conformation of the Smc5/6 holo-complex, but the proximity of the primary modification site to the α1-helix of Nse2 hints at a regulatory role; potentially one that acts to control arm ‘architecture’ (**Fig. 2**, inset *iii*) as well as performing the documented roles in promoting interactions with, and / or modulating activity of, replication fork associated-DNA helicases such as Sgs1^BLM^ (Bermudez-Lopez et al. 2016) and Mph1^FANCM^ (Xue et al. 2014).

Interestingly, SUMOylation catalysed by the Smc5/6 complex has been shown to be stimulated by the addition of single-stranded DNA (ssDNA) to reaction mixes (Varejao et al. 2018) with circular dichroism experiments showing localised changes in the secondary structure of Smc5 upon DNA-engagement. We speculate that this is likely to occur as a direct consequence of an initial binding event at the hinge-region of the complex, that leads to stimulation and auto-modification of Smc5/6, which then allows the complex to control/modulate homologous recombination activity at a stalled replication fork, presumably through regulation of specific helicases and/or other modified substrates; supported by our experimental observations in fission yeast, where mutations affecting the ability of the hinge to bind ssDNA lead to gross chromosomal rearrangements, but do not impact chromatin-binding or retention of the complex {Etheridge, 2021 #164}.

Finally, our cryo-EM structure of the apo/non-liganded state provides a highly valuable ‘stepping stone’ along the way to a fuller understanding of the set of molecular interactions and dynamics than underpin the cellular function(s) of the Smc5/6 complex. Clearly, a desirable goal is now the determination of structures for the ATP-bound and DNA-engaged states.

## MATERIAL AND METHODS

### Expression and purification

Detailed experimental procedures for both expression and purification of the *S. cerevisiae* Smc5/6 holo-complex are available in (Hallett et al. 2021). For convenience, the composition of the two buffers that have been used in this study are listed below:

**BUFFER C:** 20 mM HEPES.NaOH pH 7.5, 100 mM NaCl, 0.5 mM TCEP
**BUFFER F:** 20 mM HEPES.NaOH pH 7.5, 0.5 mM TCEP

### Cryo-EM

#### Sample preparation

Fractions eluting from a Superose 6 size exclusion chromatography column (equilibrated in BUFFER C; Cytiva Life Sciences, Little Chalfont, UK), corresponding to BS3-crosslinked holo-complex, were immediately used for grid preparation. The sample was diluted by a factor of two (with addition of BUFFER F) to reduce the overall NaCl concentration to 50 mM and yield a final concentration of 0.1 mg/ml. From this, 3 μl was applied to a freshly glow-discharged grid (Quantifoil R0.6/1 Cu 300 mesh grid; 60 seconds, 15 mA, PELCO easiGlow — Agar Scientific, Stansted, UK). Using an EM GP2 automatic plunge freezer (Leica Microsystems, Wetzlar, Germany) the grid was held in a chamber at 10 °C and 90% relative humidity, for a period of 10 seconds, before blotting for 2.5 - 4.5 seconds using the auto-sensor. The grid was immediately plunged into liquid ethane at −182 ° and then stored under liquid nitrogen until data collection.

#### Data collection

Data were collected at LonCEM (The Frances Crick Institute, London, UK) at 300 kV on a Titan Krios electron microscope (Thermo Fisher Scientific, Waltham, MA USA), equipped with a Gatan K3 detector operating in counted super-resolution mode. Movies were acquired using EPU (Thermo Fisher Scientific). Data were collected from four grids across separate sessions, without hardware binning at a calibrated pixel size of 0.55 Å. Target defocus was −1 to −3.5 μm, with a total dose of 50 electrons per Å^2^, during an exposure time of 3.9 to 5 seconds, fractionated into 38-50 frames.

#### Data processing

An overview of the workflow used to process data is provided in **Fig. S1**. In summary, all movie frames were aligned using 5 × 5 patches in Motioncor2 with dose-weighting (Zheng et al. 2017). All data were binned by a factor of 2. Contrast transfer function (CTF) parameters were estimated using CTFFIND4 (Rohou and Grigorieff 2015). Particles were picked using Topaz (Bepler et al. 2019) integrated into CryoSPARC. Initial training used a set of particles manually picked from a few micrographs, with the resultant ‘trained’ model being used to auto-pick particles across all micrographs. Apart from the ‘Head-only’ approach, all processing was carried out in parallel, using both CryoSPARC (v3.1.0, Punjani et al. 2017) and RELION (v3.1, (Zivanov et al. 2018)) software suites to yield similar set of maps. For the sake of simplicity and brevity, only the maps with the highest resolution estimates are reported here. All refinements were performed using independent data half-sets (gold-standard refinement) and resolutions determined based on the Fourier shell correlation (FSC = 0.143) criterion (Rosenthal and Henderson 2003).

#### Holo-complex

After several rounds of 2D classification in cryoSPARC, particles from ‘good’ classes were used for an initial *ab initio* reconstruction. The resultant particles were exported to Relion and then re-classified in 2D. Subsequent rounds of 3D classification in Relion yielded 6 classes, of which one (1: HOLO-COMPLEXl; 17,152 particles, representing 13% of the input) provided the highest resolution after 3D refinement with a soft-edged mask and solvent flattening. Post-processing produced a map at 10.8 Å resolution.

#### Hinge/Arm and Head

Working in parallel, using cryoSPARC, particle subtraction was used to remove the ‘head-end’ of the complex, before additional rounds of 3D classification, refinement and post-processing yielded a map at 8.53 Å (2: HINGE/ARM; 106,660 particles, 62.4% of input). A similar strategy was used to remove the ‘hinge-end’ of the complex, however further rounds of processing was halted due to the superior results obtained via the ‘head-only’ approach described below.

#### Head-only

A second trained ‘picking model’ was used to identify particles corresponding to just the ‘head-end’ of the complex. After several rounds of 2D classification, particles from ‘good’ classes were used for an initial *ab initio* reconstruction. After iterative rounds of 3D classification and refinement, one class was selected to take forward into non-uniform refinement, after removal of density corresponding to the HALO-tag attached to the Nse4 subunit by particle subtraction (3: HEAD-ONLY; 84,810 particles, 51.7% of input). Post-processing yielded a map at 6.5 Å resolution.

#### Composite Map and Model Building

An initial pseudo-atomic model for the Smc5/6-complex was generated by fitting of Phyre2-generated homology models (Kelley et al. 2015) into map segments using programs of either the PHENIX software suite (Liebschner et al. 2019) or ChimeraX (Pettersen et al. 2021; Afonine et al. 2018; Liebschner et al. 2019), with additional manual positioning in either Coot (Emsley et al. 2010) or PyMOL (The PyMOL Molecular Graphics System, Version 2.3.2, Schrödinger LLC). Phenix.combine_focused_maps was used to generate a composite map, using each of the reported maps with the initial model as an alignment reference. The initial homology models were subsequently replaced with those generated by AlphaFold upon their public release. The fit of the overall model was optimised by real-space refinement in PHENIX against the composite map.

#### Data availability

The two maps used to generate the composite cryo-EM volume have been deposited in the Electron Microscopy Data Bank (EMDB) with accession codes EMD-13893 (head-end of complex) and EMD-13894 (hinge and arm-region). Real-space refined coordinates for the Smc5/6 model have been deposited in the Protein Data Bank (PDB) with accession code PDB-7QCD. The accompanying composite cryo-EM volume has been deposited with accession code: EMD-13895.

### Experiments in yeast

Yeast strains were generated using the following procedure. Synthetic DNA encoding the genomic sequence for both WT and mutant versions of each gene were purchased as ‘Strings DNA Fragments’ from GeneArt (Thermo Fisher Scientfic, UK). These were cloned into the vector pAW8-natMX6 (Watson et al. 2008), at the PaeI / SalI restriction sites, through Gibson assembly via short regions of homology included during synthesis. PCR was then used to amplify the gene and associated nourseothricin resistance module for introduction into the endogenous locus of diploid yeast cells, using lithium acetate transformation.

#### Yeast Strains

Strains were generated from 3 individual haploid isolates, generated from tetrad dissections.

| Strain # | Genotype | Reference |
| --- | --- | --- |
| 2655 | *MATalpha [leu2-3 trp1-1 ura3-1 can1-100 ADE2 his3-11,15]; haploid isolate, derived from W303* | (Thomas and Rothstein 1989) |
| 2747<br>diploid | *MATa/alpha [leu2-3 trp1-1 ura3-1 can1-100 ade2-1/ADE2 his3-11,15 SMC5/smc5-Y961A, W964A]* | This study |
| 2753<br>isolate 1 | *MATa [leu2-3 trp1-1 ura3-1 can1-100 ADE2 his3-11,15 smc5-Y961A, W964A]* | This study |
| 2755<br>isolate 2 | *MATalpha [leu2-3 trp1-1 ura3-1 can1-100 ADE2 his3-11,15 smc5-Y961A, W964A]* | This study |
| 2756<br>isolate 3 | *MATalpha [leu2-3 trp1-1 ura3-1 can1-100 ADE2 his3-11,15 smc5-Y961A, W964A]* | This study |
| 2745<br>diploid | *MATa/alpha [leu2-3 trp1-1 ura3-1 can1-100 ade2-1/ADE2 his3-11,15 SMC5/smc5-F972A, L978D, L981N]* | This study |
| 2793<br>diploid | *MATa/alpha [leu2-3 trp1-1 ura3-1 can1-100 ade2-1/ADE2 his3-111,15 NSE1/nse1-F217A, E228R, R242A]* | This study |
| 2794<br>isolate 1 | *MATa [leu2-3 trp1-1 ura3-1 can1-100 ADE2 his3-111,15 nse1-F217A, E228R, R242A]* | This study |
| 2795<br>isolate 2 | *MATa [leu2-3 trp1-1 ura3-1 can1-100 ADE2 his3-111,15 nse1-F217A, E228R, R242A]* | This study |
| 2796<br>isolate 3 | *MATalpha [leu2-3 trp1-1 ura3-1 can1-100 ADE2 his3-111,15 nse1-F217A, E228R, R242A]* | This study |

### Figures

Molecular images were generated using either PyMOL (v. 2.3.2) or ChimeraX (v. 1.1.1).

## ACKNOWLEDGEMENTS

The authors would like to thank Dr. Adam Watson, Dr. Alessandro Bianchi, Prof. Uli Rass, Prof. Antony Carr [AMC] and Prof. Laurence Pearl [LHP] (University of Sussex) for advice, discussion, and access to reagents/equipment. We are also grateful to the cryo-EM facility at the University of Sussex (funded by a Wellcome Trust award enhancement grant 095605/Z/11/A to LHP and the RM Phillips Trust). The work was supported by funding from: University of Sussex Strategic Development Fund [JB], Wellcome [110047/Z/15/Z; AMC], Cancer Research UK [C302/A24386; AWO and LHP] and MRC [MR/PO18955/1; JMM and AWO]. Funding for open access charge: Medical Research Council, University of Sussex Open Access Team (Library).

---

**Supplementary Figure 1.**
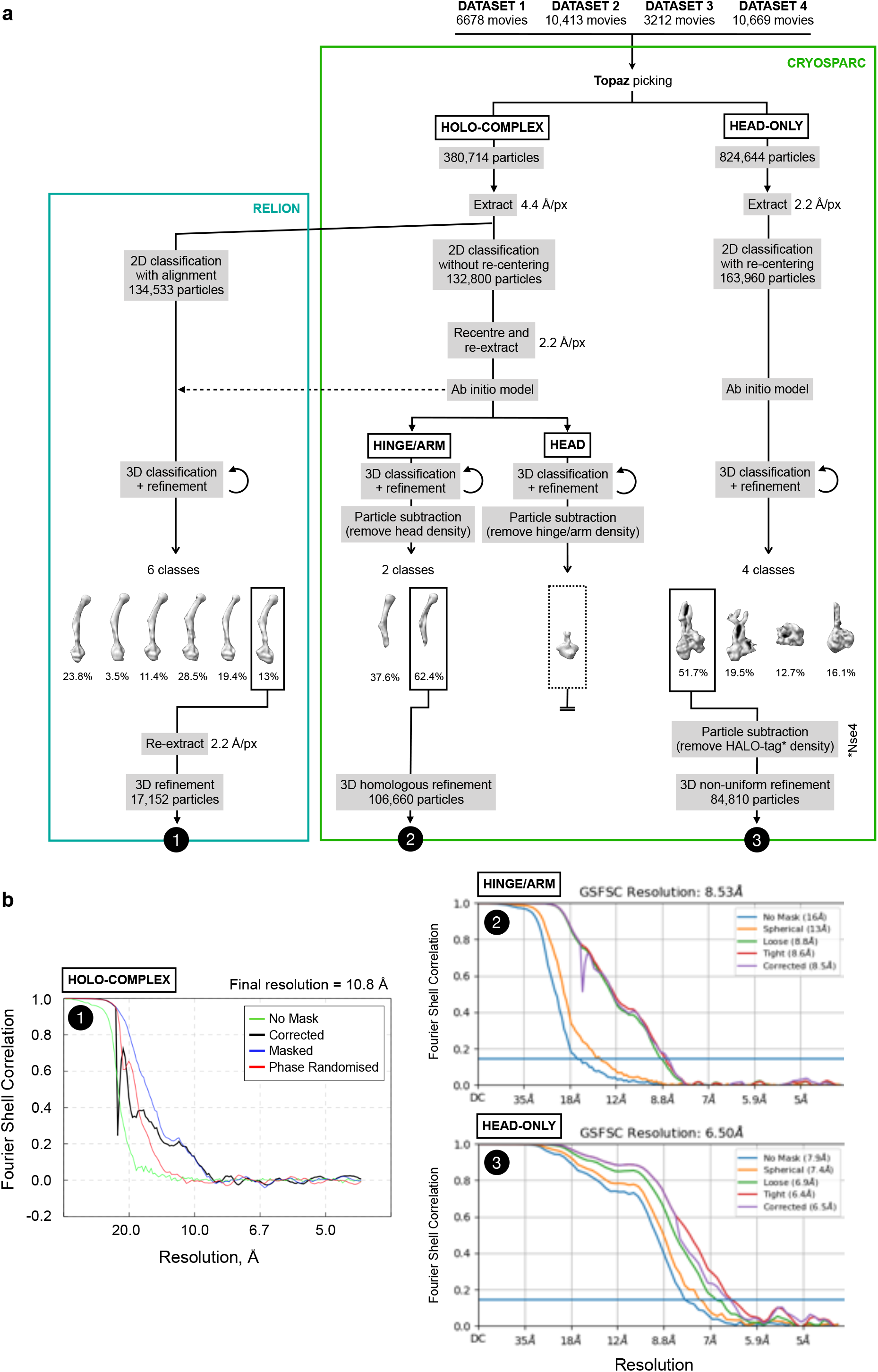
(a) Summary of image-processing workflow, used to determine the structure of the budding yeast Smc5/6 holo-complex. (b) Fourier Shell Correlation (FSC) curves estimating the average resolution of the indicated maps.

**Supplementary Figure 2.**
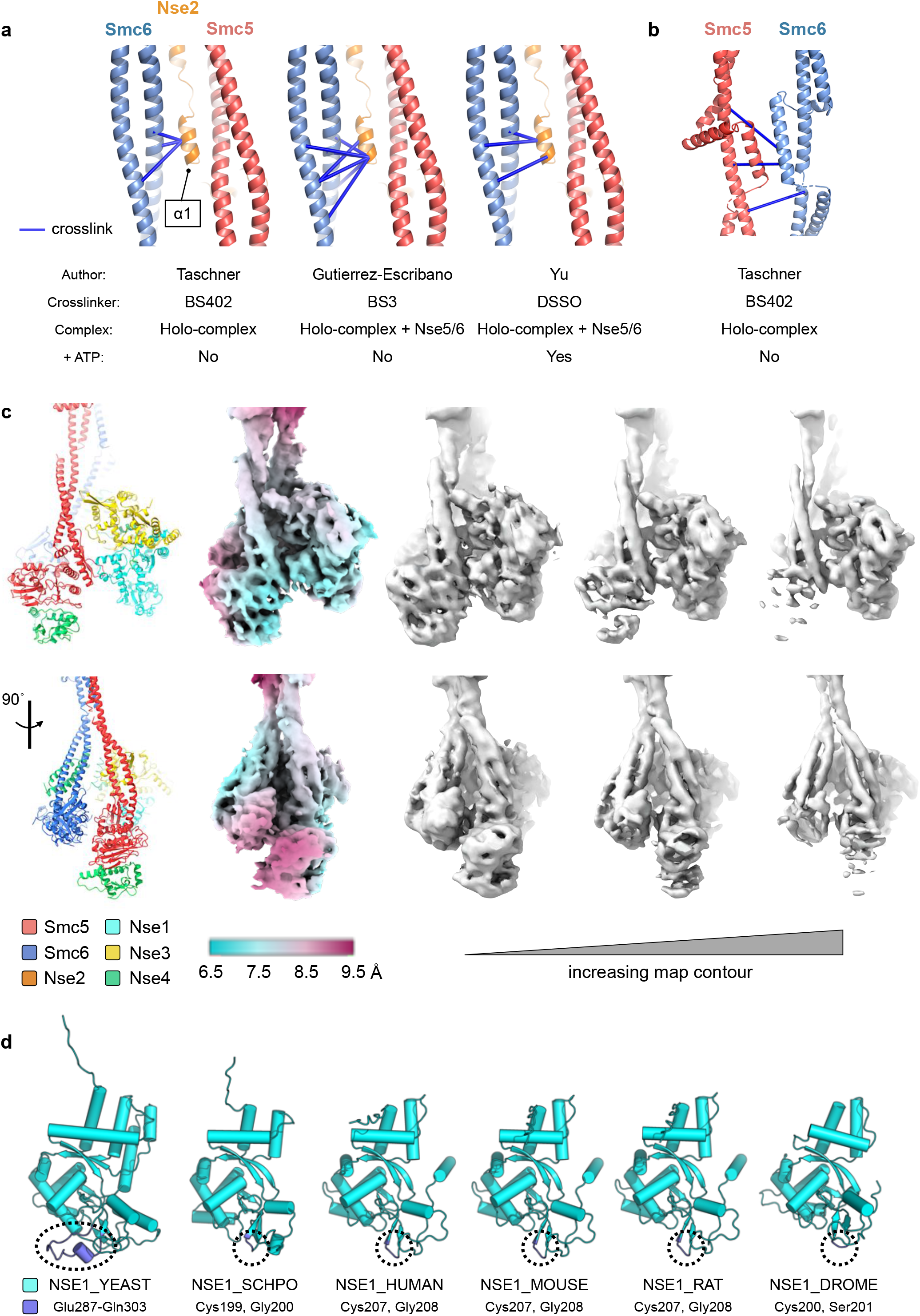
(a) Selected cross-linking mass spectrometry (XL-MS) data taken from 3 separate studies, supporting interaction of the first helix of Nse2 (α2) with the arm of Smc5. Study, type of crosslinker used, complex composition and nucleotide status are summarised below the molecular graphics. Cross-links were visualised using the PyXLinkViewer plugin (Serrano et al. 2020) for PyMOL (The PyMOL Molecular Graphics System, Version 2.3.2, Schrödinger, LLC). XL-MS data reported in (Taschner et al. 2021), (Gutierrez-Escribano et al. 2020), (Yu et al. 2021). (b) Selected XL-MS data taken from (Taschner et al. 2021) supporting interaction of the two SMC ‘joints’ in the Smc5/6 holo-complex. (c) Different representations for the ‘head-end’ of the Smc5/6 complex. Sequentially from left to right: secondary structure molecular cartoon; estimate of local resolution as calculated by ResMap-1.1.4 (Kucukelbir, Sigworth, and Tagare 2014) mapped onto the surface of the composite cryo-EM map; composite map shown at increasing contour levels. See associated keys for additional details. (d) AlphaFold predicts the presence of a budding yeast-specific loop insertion in Nse1 (amino acids 287-303, coloured dark blue).

**Supplementary Figure 3.**
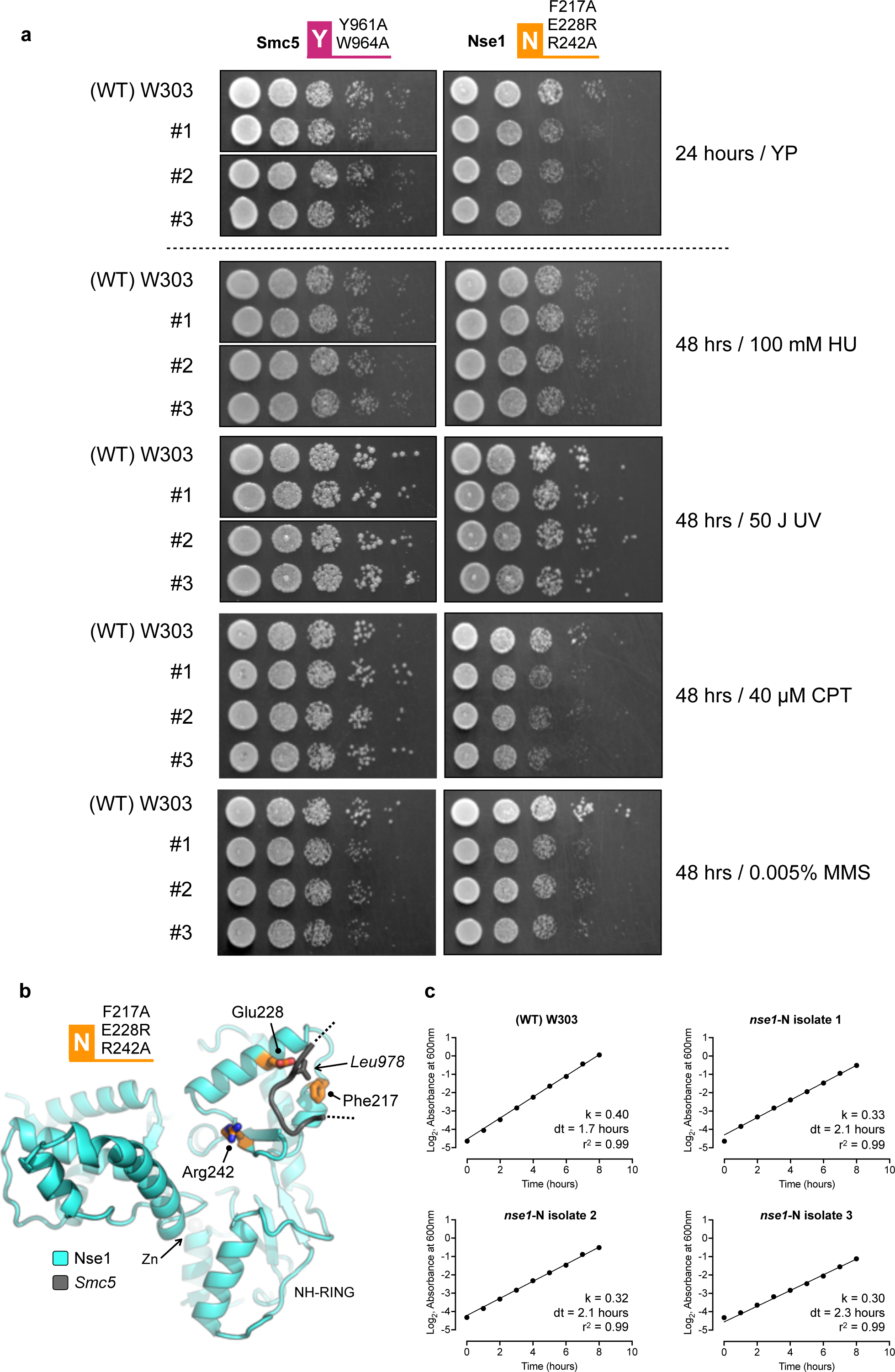
(a) Yeast strains carrying the Y-set of Smc5 mutations are not sensitive to a range of genotoxic agents and have no obvious growth defect. Those carrying the N-set of Nse1 mutations are mildly sensitive to treatment with camptothecin (CPT) or methyl methanesulphonate (MMS). Dose, type of treatment and duration of growth are as indicated; where HU = hydroxyurea, UV = ultraviolet light. The W303 strain is included as a wild-type (WT) control. (b) Molecular secondary structure cartoon highlighting the position of the mutated amino acid residues in Nse1 (carbon atoms coloured in orange and shown in ‘stick’ representation) relative to that of the Smc5 ‘W-loop’ (coloured in dark grey, with the side chain of Leu978 shown in ‘stick’ representation). Please also see associated key for additional details. (c) Representative growth curves for the 3 isolates containing the N-set of mutations introduced into Nse1. The W303 strain is included as a wild-type (WT) control. Data are fitted to an Exponential (Malthusian) grown curve, with k defining a rate constant (slope), dt indicating the calculated doubling time, and r^2^ a goodness of fit parameter.

